# Distinct host preconditioning regimens differentially impact the antitumor potency of adoptively transferred Th17 cells

**DOI:** 10.1101/2023.12.18.572179

**Authors:** Megen C. Wittling, Hannah M. Knochelmann, Megan M. Wyatt, Guillermo O. Rangel Rivera, Anna C. Cole, Gregory B. Lesinski, Chrystal M. Paulos

## Abstract

**Background:** Mechanisms by which distinct methods of host preconditioning impact the efficacy of adoptively transferred antitumor T helper cells is unknown.

**Methods:** CD4^+^ T cells with a transgenic TCR that recognize TRP-1 melanoma antigen were polarized to the T helper 17 (Th17) phenotype and then transferred into melanoma-bearing mice preconditioned with either total body irradiation or chemotherapy.

**Results:** We found that preconditioning mice with a non-myeloablative dose of total body irradiation (TBI of 5 Gy) was more effective than using an equivalently dosed non-myeloablative chemotherapy (CTX at 200 mg/kg) at augmenting therapeutic activity of anti-tumor TRP-1 Th17 cells. Anti-tumor Th17 cells engrafted better following preconditioning with TBI and regressed large established melanoma in all animals. Conversely, only half of mice survived long-term when preconditioned with CTX and infused with anti-melanoma Th17 cells. IL-17 and IFN-g produced by the infused Th17 cells, were detected in animals given either TBI or CTX preconditioning. Interestingly, inflammatory cytokines (G-CSF, IL-6, MCP-1, IL-5, and KC) were significantly elevated in the serum of mice preconditioned with TBI versus CTX after Th17 therapy.

**Conclusions:** Our results indicate, for the first time, that the antitumor response, persistence, and cytokine profiles resulting from Th17 therapy are impacted by the specific regimen of host preconditioning. This work is important for understanding mechanisms that promote long-lived responses by ACT, particularly as CD4^+^ based T cell therapies are now emerging in the clinic.

## Background

Adoptive T cell transfer (ACT) therapy involves the infusion of tumor-specific T cells into patients, and ranges from tumor infiltrating lymphocytes (TIL) to lymphocytes engineered with antigen receptors (CAR or TCR) [1–6]. Optimizing ACT to treat patients with aggressive malignancies is an important undertaking, given that this treatment shows promise in the clinic [7–10]. In particular, lymphodepleting preconditioning regimens prior to ACT are standard modalities to enhance the potency of ACT and do so via multiple mechanisms [11–14]. Yet, it remains unknown what type of host preconditioning is most ideal for CD4^+^ T helper cell therapies, a lymphocyte population that is emerging as promising to treat patients [15, 16]. To date, how different forms of non-myeloablative preconditioning regimens impact the efficacy of transferred antitumor T helper cells has not been explored in a systematic manner.

Previous research has shown that lymphodepletion via TBI or cyclophosphamide is an effective regimen to enhance the anti-tumor activity of transferred anti-melanoma CD8^+^ T cells [17]. Lymphodepletion transiently ablates immunosuppressive host cells, such as regulatory T cells and myeloid derived suppressor cells, which actively blunt CD8^+^ T cell functionality [18–20]. Both forms of lymphodepletion create “space” via depleting host NK cells and lymphocytes, in turn inducing homeostatic cytokines that can be used to instead support the transferred T cells [21, 22]. Additionally, TBI mediates the systemic translocation of microbial ligands from the gut, known to activate antigen presenting cells, in turn enhancing the function of transferred CD8^+^ TIL [14]. Radiation can also activate DCs by mediating the systemic release of damage-associated molecular patterns (DAMPs) and high-mobility group box 1 protein (HMGB1), resulting in improved adaptive responses to tumors [23]. Cyclophosphamide has specific benefits as well – increasing IFNa production and aiding Th17 differentiation [24, 25]. Most recently, both Th17 cells and hybrid Th1/17 cells (able to co-secrete IL-17A and IFN-g) were enriched in patients with breast cancer responsive to neoadjuvant chemotherapy [26]. However, despite these known benefits of lymphodepletion, the effects of these regimens on antitumor Th17 therapy are unknown.

Non-myeloablative total body irradiation (TBI) is a common method to mediate host lymphodepletion in mice prior to ACT therapy [17, 27–31]. Aligned with this pre-clinical approach, a series of clinical trials at the NCI were conducted using TIL in patients and giving them various regimens of total body irradiation [9]. Yet, most individuals given ACT are preconditioned with nonmyeloablative chemotherapies, including cyclophosphamide (CTX) and/or fludarabine (FLU), as TBI at a dose of 1200 cGy in humans has been associated with toxicities including more profound neutropenia and thrombotic microangiopathy [17, 32]. In mice, it was reported that escalating the intensity of lymphodepletion by increasing doses of fractionated TBI stepwise elevated features of toxicity in animals, based on heightened cytokine storm signatures and translocation of gut microbes [33]. While this approach improved outcomes in the preclinical setting, it was not without undue side effects. This work points to focusing on using non-myeloablative preparative regimens, including CTX (*which is more common in the clinic*) to augment T helper therapy. However, the mechanistic understanding of how these various preconditioning regimens impacts infused cells and their therapeutic efficacy is unclear. Further, we posit that these approaches of lymphodepletion are not interchangeable and harbor distinct mechanistic differences that may impact subsequent ACT *in vivo*. A greater appreciation for these differences may inform the field for optimizing pre-clinical studies of CD4^+^ T cell-based ACT in murine tumor models.

Overarchingly, there are many advantageous aspects of preconditioning that remain to be investigated in the context of CD4^+^ cellular cell therapies. Clinical trial data supports the use of chemotherapy prior to T cell infusion, with CTX administration leading to both increased T cell engraftment and persistence in patients [34, 35]. Given that both CTX and TBI allow for improved engraftment and deplete immunosuppressive host cells, we sought to define how these two distinct methods of lymphodepletion impact the therapeutic efficacy anti-melanoma Th17 cells, which is emerging as promising in the field. Herein we examine how these two forms of host preconditioning impact the efficacy of a novel adoptive Th17 cell therapy that our team and others have found are potent against tumors [36–42].

## Results

### Th17 cells mediate superior survival protection in mice compared to Th1 cells

To test the potency of antitumor CD4^+^ Th1 versus Th17 cells as an adoptive immunotherapy, we used the transgenic TRP-1 model where CD4^+^ T cells express a TCR that recognizes tyrosinase-related peptide (TRP) on melanoma [38]. Mice bearing B16F10 melanoma were preconditioned with lymphodepletion using 5Gy total body irradiation (TBI) and then were treated with infusion of TRP-1 Th1 or Th17-cytokine programmed lymphocytes (Fig. 1A). As expected, TRP-1 Th1 cells secreted more IFN-g and nominal levels of IL-17A *ex vivo* upon reactivation with their cognate antigen (Figures 1B-C). Conversely, Th17 cells mainly produced IL-17A, but little IFN-g after programming *ex vivo* (Figures 1B-C). Moreover, the infusion of TRP-1 Th17 cells mediated the most efficacious anti-tumor responses in mice, as denoted by the ability of 22 of 26 animals (∼85%) to survive nearly two months post ACT therapy with Th17 versus Th1 therapy (Figure 1D) [38]. These results corroborate those from other laboratories who have also found that Th17 cell therapy is effective against various solid tumors, often mediating long-lived responses in mice [38–43].

**Figure 1:**
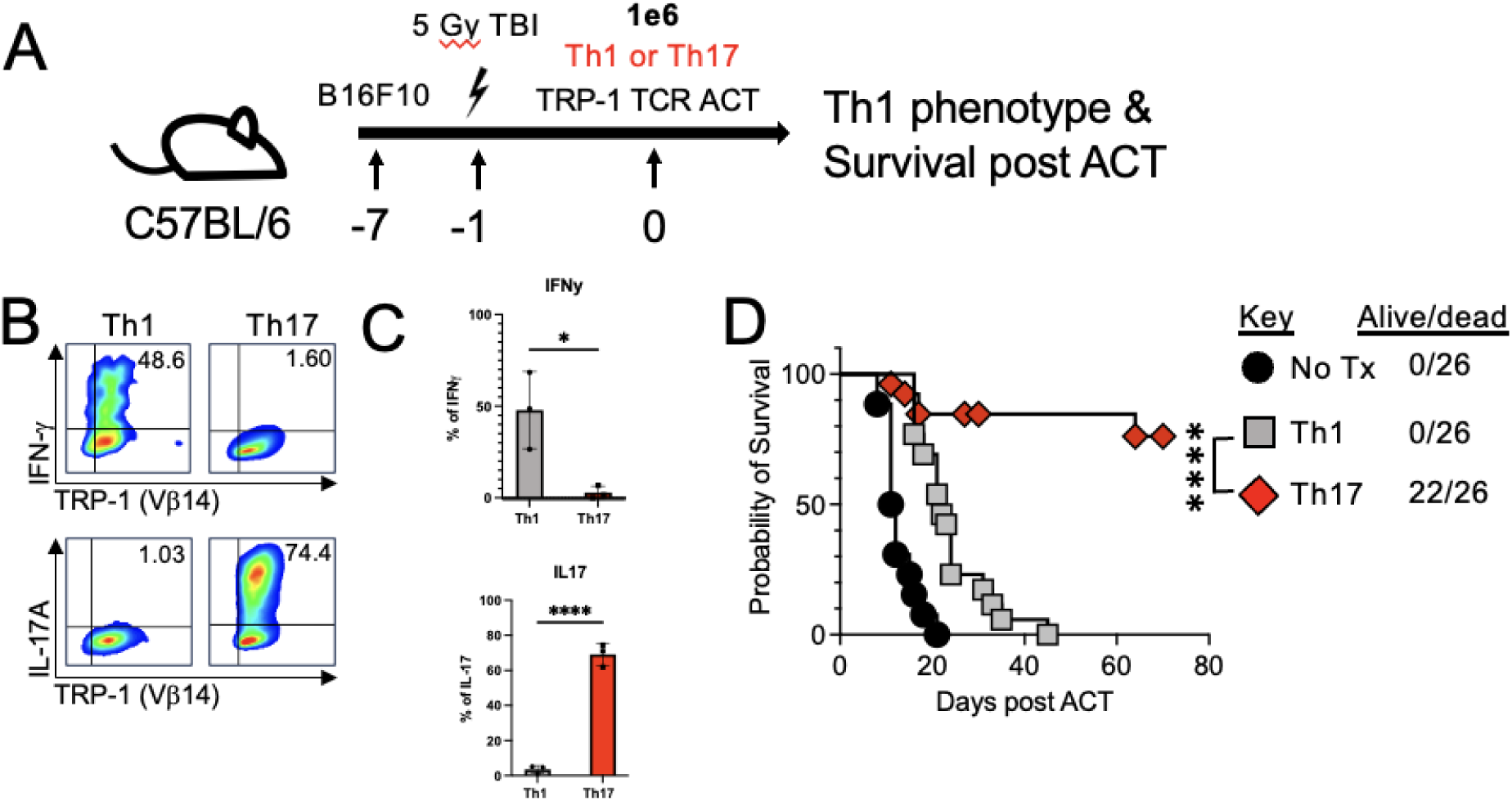
Adoptive transfer of TRP-1 Th17 cells significantly improves survival of mice bearing established melanoma, as compared to transferred Th1 cells. (A) Mouse experimental model. C57BL/6 mice bearing B16F10 melanoma were preconditioned with 5 Gy total body irradiation one day prior to adoptive cell transfer of TRP-1 specific Th1 (IL-12, aIL-4, IL-2) or Th17 (IL-6, IL-21, IL1b, TGF-b, aIFN-g, aIL-4, aIL-2) cells (1e6). (B) Representative flow plots and (C) Bar graph of IFN-LJ and IL-17A production by TRP-1 subsets post reactivation with PMA/ionomycin on day 7 after expansion (n=3). Analysis via t-test. (D) Survival curve from three combined experiments of mice (n = 6-10 mice/group) with day 7 established melanoma. Th17 vs Th1 P<0.0001 Log-rank (Mantel-Cox) test.

### TBI as lymphodepletion prior to Th17 infusion improves antitumor activity in mice

Given the potency of our antigen specific Th17 cells, we hypothesized that preconditioning may not be needed for its efficacy. We explored this idea via comparing two distinct preconditioning regimens: chemotherapy or total body irradiation prior to ACT. This cell therapy experiment was done by comparing a chemotherapy to TBI, or no preconditioning prior to transfer of Th17 cells into mice with melanoma (Figure 2A). Moreover, we tested the impact of different preconditioning methods of lymphodepletion on *in vivo* tumor growth and survival of mice.

**Figure 2:**
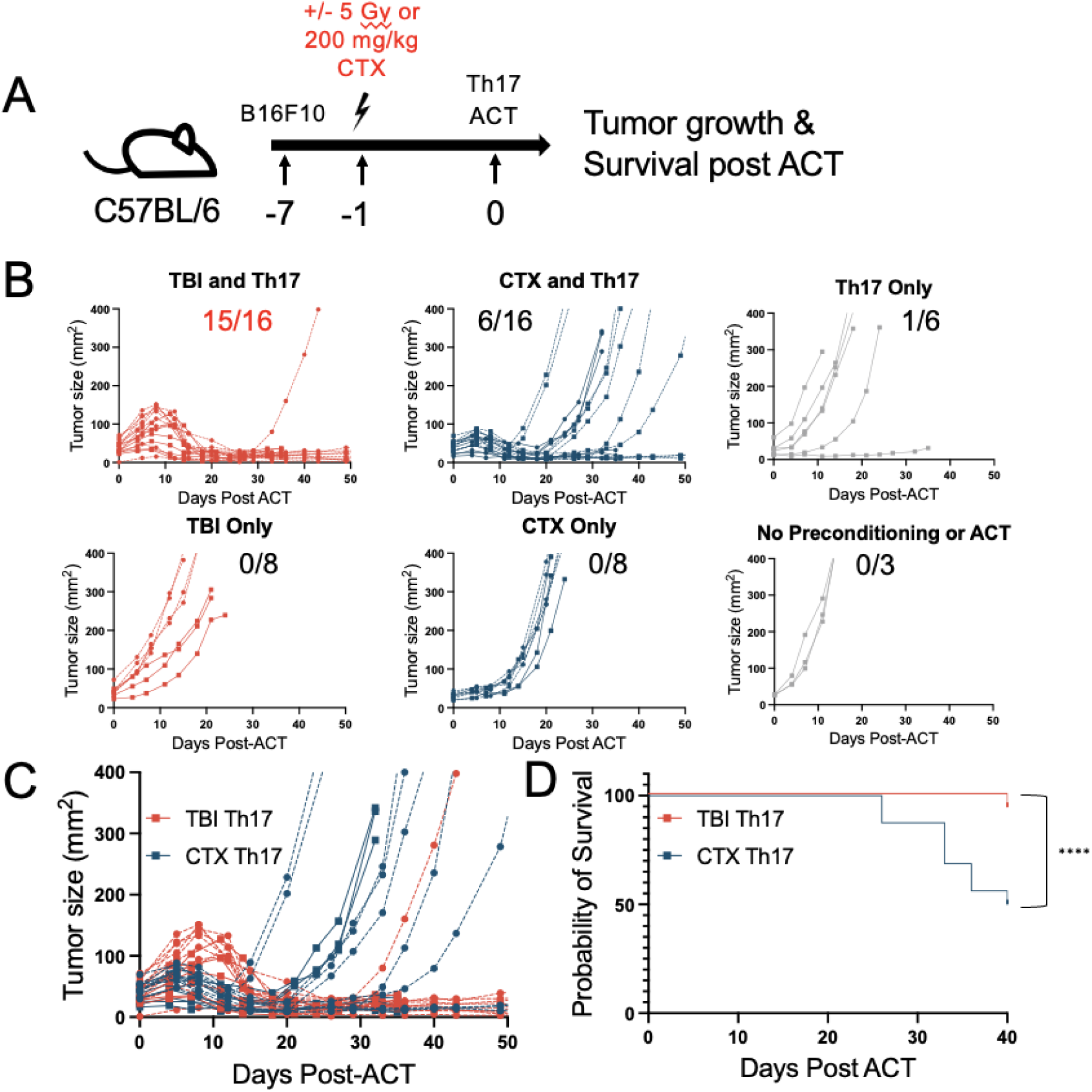
Th17 Therapy mediated superior survival in melanoma-bearing mice conditioned with TBI compared to unconditioned mice or mice preconditioned with CTX. (A) Mouse experimental model. C57BL/6 mice bearing B16F10 melanoma were preconditioned with 5 Gy total body irradiation, 200 mg/kg CTX, or no preconditioning one day prior to adoptive cell transfer of TRP-1 specific Th17 cells. (B) Tumor growth curves (n=2). Note that the dotted lines are from one independent experiment and solid lines are from a second independent experiment. Tumors were measured as length x width. (C) Overlay of TBI preconditioned and CTX preconditioned mice given ACT for two independent experiments (n=32 mice). (D) Survival curve for each TBI+ACT and CTX+ACT groups. Log-rank test comparison of the two groups resulted in p<0.0001.

In contrast to our hypothesis that preconditioning is dispensable for the antitumor effects of our potent Th17 ACT, we found that if mice were not preconditioned with some form of a lymphodepletion, then all but one animal had rapidly growing tumors and reached tumor endpoint within a month (Figure 2B). Radiation or chemotherapy alone (without infusion of Th17 cells) was not effective. However, when mice were pretreated with either CTX or TBI prior to TRP-1 Th17 therapy, mice in both cohorts achieved cures, albeit with different efficacies (Figures 2B). For example, most animals given TBI (15/16 mice) experienced tumor regression (Figure 2B) and curative responses when infused with antitumor Th17 cells. In contrast, CTX was less efficacious overall, as only a few mice were cured via ACT, with most (10/16) succumbing to disease between 2-4 weeks post infusion of Th17 cells. This stark contrast in response can be appreciated in the overlay of these two conditions as shown in Figure 2C. The two lymphodepletion regimens differentially impacted survival as well (Figure 2D), as evidenced by improved outcomes in mice receiving Th17 ACT that were preconditioned with TBI versus CTX (p<0.0001).

### A unique cytokine signature is induced in the serum of mice treated with TBI Th17 therapy

Because TBI preconditioning of mice was more effective than CTX when treating with Th17 ACT, we next examined what factors that might potentially differentiate these therapies. We hypothesized the induction of a cytokine profile may be associated with improved efficacy. We therefore assessed levels of serum cytokines in these mice 10 days after ACT. IFN-g and IL-17 (produced by transferred Th17 cells) were elevated in animals given either lymphodepletion method when infused with Th17 cells. Note, these two cytokines were only elevated in mice treated with a preconditioning regimen combined with Th17 ACT (Figure 3A). In CTX preconditioned mice with ACT, there are some mice with high IL-17 expression (∼50 pg/mL) and half with lower IL-17 (∼5 pg/mL) (Figure 3A). Interestingly, the mice in this first cohort with more serum IL-17 survived longer (nearly two months post ACT) when given CTX. Overall, these findings demonstrate that preconditioning increased functionality of transferred Th17 cells.

**Figure 3:**
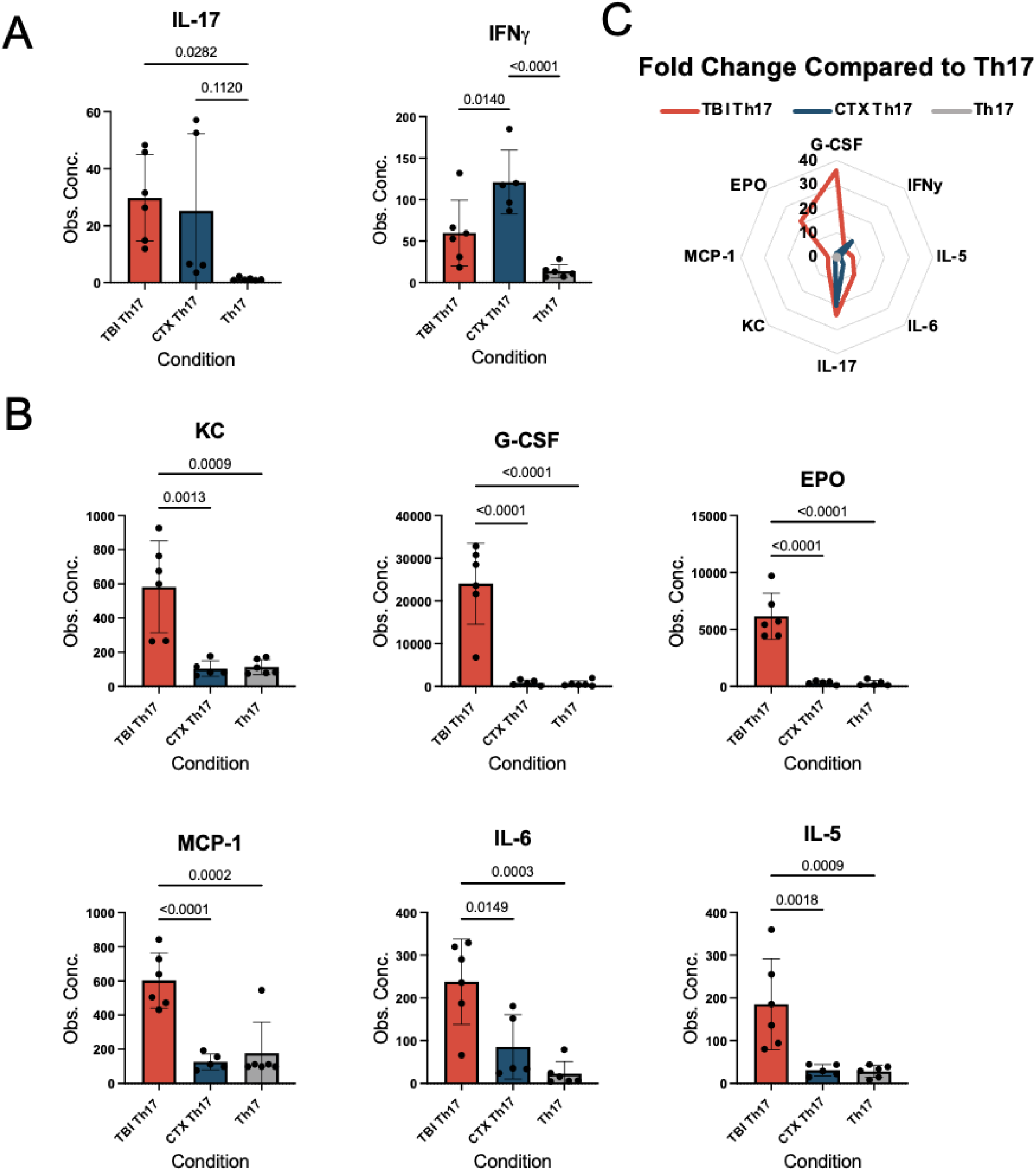
TBI potentiates unique inflammatory but not Type 1 or Type 17 cytokines in mice compared to CTX conditioning. (A) Observed concentration in pg/mL of cytokines MCP-1, IL-6, IL-5, KC, and G-CSF in serum collected from mice. Statistics calculated via one-way ANOVA. (B) Observed concentration in pg/mL of IL-17 and IFNg. Statistics calculated via one-way ANOVA. (C) Radar Plot displaying the fold change of TBI Th17 and CTX Th17 groups compared to Th17 alone control.

Differences in cytokine profiles in mice given CTX versus TBI prior to Th17 therapy was observed. Keratinocyte chemoattractant (KC), granulocyte colony-stimulating factor (G-CSF), monocyte chemoattractant protein-1 (MCP-1), EPO, IL-6, and IL-5 were vastly elevated in the serum of mice treated with Th17 cells and conditioned with TBI compared those given CTX (Figure. 3B). Fold change increase in these cytokines is visualized in Figure 3C, with a systemic cytokine induction ranging for 3-36X more in animals given TBI versus nonpreconditioned mice.

### Infused Th17 cells persist best in mice preconditioned with TBI

Another parameter used as a biomarker for successful ACT therapy is the degree to which the infused T cell products engraft and persist in patients. Compared to mice only given Th17 therapy alone (without preconditioning), both lymphodepletion methods permitted comparable engraftment of transferred donor CD4^+^ cells (Figure 4A). Yet, because Th17 cells were proliferating to a profoundly higher degree in mice given TBI compared to CTX at Day 7 (Figure 4B), we hypothesized that donor Th17 cells may engraft and subsequently persist better in animals given TBI versus CTX. Indeed, more than one month post ACT (39 days later), more TRP-1 Th17 cells were detected across multiple different tissues if the animals were preconditioned with TBI compared to CTX (Figure 4C). These data suggest the TBI and CTX differentially impact Th17 functionality and persistence, as opposed to the acute abilities of these cells to engraft. Our findings also underscore that either form of lymphodepletion improves success of Th17 therapy in this well-established melanoma model system.

**Figure 4:**
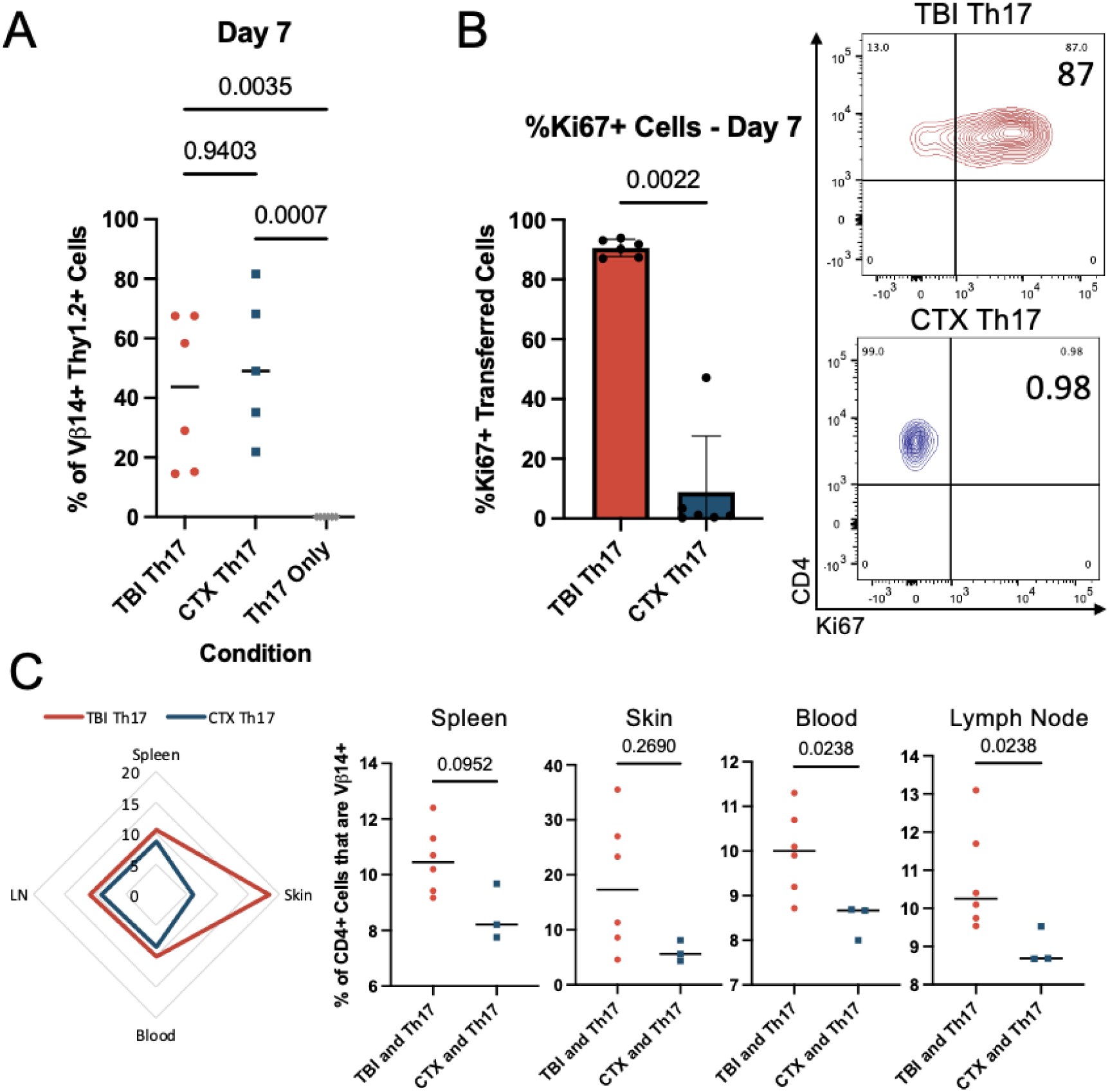
Adoptively Transferred Th17 cells prevail best when mice are preconditioned with Total Body Irradiation. A. Th17 cells from ACT identified via Vb14 and Thy1.2 positivity via flow. Gated on live/CD3+/CD4+. Percent of donor cells in the CD4 compartment of the peripheral blood on Day 7 in mice with 5 Gy TBI, CTX, or no preconditioning. Analysis via Mann-Whitney Test. B. Peripheral blood taken from mice on Day 7 after ACT. Gated on live/CD3+/CD4+/Vb14+/Thy1.2+. Increased staining of proliferation marker Ki67 in adoptively transferred Th17 cells with TBI preconditioning compared with CTX. C. Persistence at endpoint (D39) identified via Vb14+ cells gated on live/CD3+/CD4+ in the peripheral blood, lymph node, skin, and spleen.

**Figure 5:**
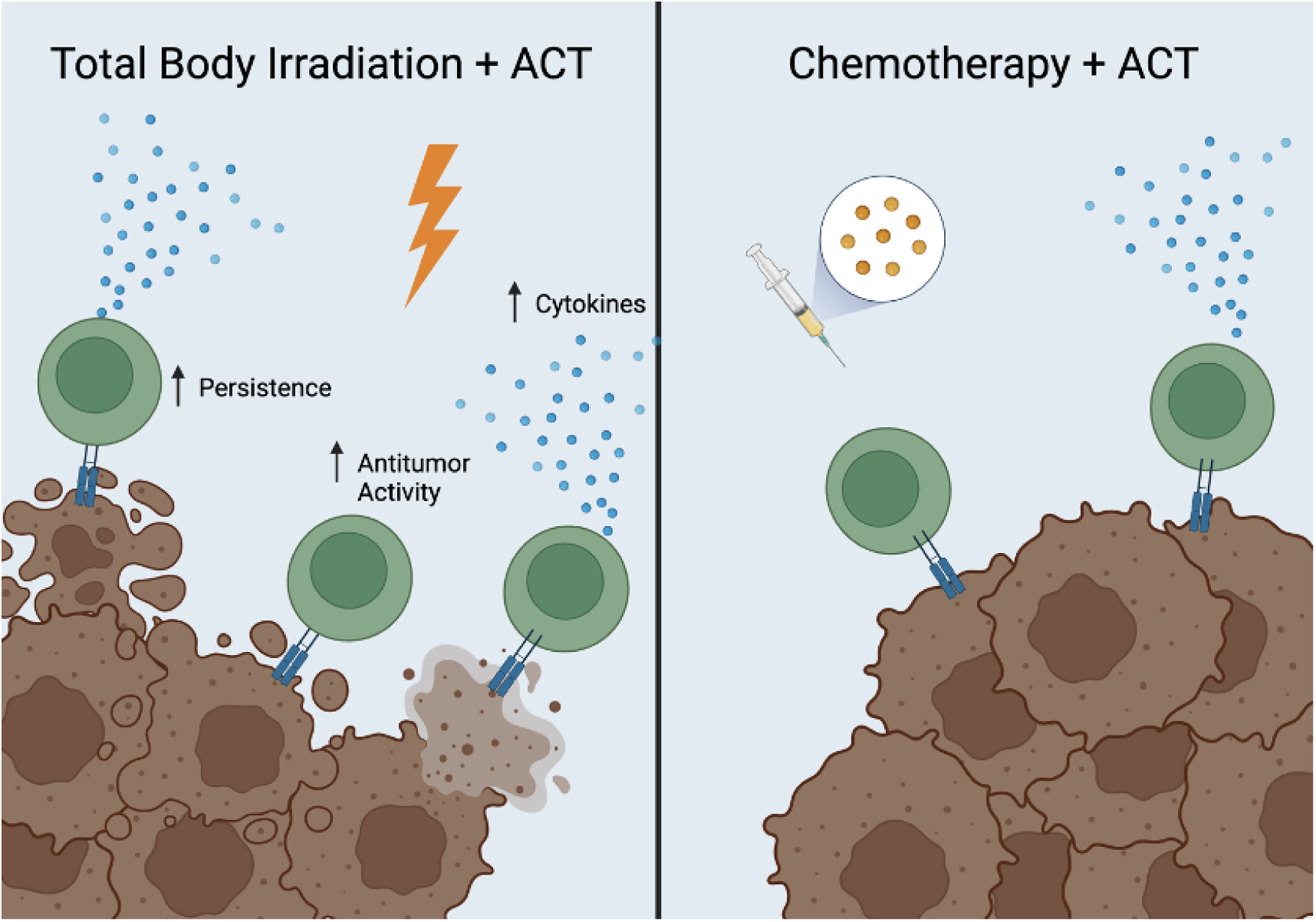
Summary Figure. CD4 Th17 ACT has superior efficacy when preconditioning with total body irradiation compared to chemotherapy is used. There is differential induction of cytokines, enhanced survival, and increased persistence of adoptively transferred cells when using this method. Finally, chemotherapy alone is superior to no preconditioning for ACT, but results in decreased survival of mice and decreased amounts of cytokines compared to mice receiving TBI. Made using BioRender.

## Discussion

Cellular therapy has revolutionized medicine for patients [2, 44, 45]. Yet ACT therapy is less effective in treating patients with solid tumors for several reasons, including 1) difficulty in T cell trafficking to the tumor, 2) decreased T cell fitness, and 3) immunosuppression in the tumor microenvironment (TME) [2, 46–48]. Regarding the TME, regulatory T cells, fibroblasts, and myeloid derived suppressor cells (MDSCs), blunt T cell survival [49]. However, despite challenges faced in the treatment of solid tumors with cellular therapy, one key solution to this problem may be to defining the way to precondition the patient prior to ACT therapy.

Fundamentally, there is a great need to understand how transferred T cells interact with the tumor and with host immune cells. In essence, this knowledge could improve our application of preconditioning to maximize ACT efficacy, including Th17-based cell products. A wealth of data has been reported on how manipulating the host via lymphodepletion can augment infused T cells [19, 22, 50]. Lymphodepletion augments ACT therapy in a multifactorial fashion – depleting host Tregs and MDSCs, stimulating the innate immune system, and enhancing T cell activation [14, 50]. Finally, lymphodepletion reduces host cells that consume IL-7 and IL-15, in turn empowering the transferred T cells to consume these them instead [22]. Yet it is unknown how host lymphodepletion impacts the effectiveness of antitumor CD4^+^ T helper cells, which have emerged in the clinic as a promising therapy[51–53]..

We discovered that either form of lymphodepletion (TBI or CTX) improved antitumor Th17 therapy. Albeit lymphodepletion with TBI best augmented Th17 therapy, resulting in more cures in animals. Tumor-bearing animals who were untreated died quickly, underscoring that host preconditioning only slightly delays tumor growth and that Th17 therapy is needed to mediate durable immunity. Several clues on the cytokines that were distinctly present in animals given TBI + Th17 might inform the mechanisms for why this therapy might have been more effective. Key cytokines that regulate neutrophils (KC, G-CSF), mature eosinophils (IL-5), prevent apoptosis of red blood cells (EPO) and mediate a “cytokine storm” (MCP-1, IL-6), were heightened in animals given TBI + Th17 therapy versus CTX + Th17 therapy [54–57]. Future studies could test if administering these cytokines to animals given CTX could augment antitumor Th17 therapy to the degree seen in animals given TBI + Th17 therapy.

We consistently found that infused TRP-1 Th17 cells persisted better in multiple organs of animals that were given TBI compared to CTX, and this factor might explain why this therapy was more effective. Future studies to elaborate on the specific host cells present in mice given either TBI or CTX are needed to better define how these host elements might regulate transferred Th17 cells. Collectively, this new body of work provides deeper insight into host responses to Th17 therapy, implying novel mechanisms that might promote effective ACT. For example, we detect resident memory Th17 cells in animals given TBI therapy, which might play a role in producing protective immunity [31]. Perhaps IL-6 contributes to the durable memory of infused Th17 cells, as we reported that blocking it impaired antitumor efficacy of this ACT approach [31]. Neutrophils can also play a positive role in TRP-1 CD4 T cell therapy [58]. As KC and G-CSF induce neutrophils, they might be responsible for augmenting the TBI + Th17 therapy. Future studies in our lab will test this idea.

While preconditioning mice with TBI prior to Th17 therapy was more potent that conditioning with TBI, we are not advocating the use of TBI in human patients given cellular therapy. Rather, our work reveal that it is important to further disentangle the mechanisms by which TBI mediates robust antitumor responses and use this insight to improve next generation ACT protocols. CTX is used frequently in patients as a means of preconditioning for several important reasons: TBI can have many detrimental side effects and has only been conducted comprehensively in one investigation (the Surgery Branch NCI) in melanoma patients treated with autologous TIL products [9, 32, 59]. Also, this study, when comparing CTX and TBI in a melanoma TIL ACT setting, found that chemotherapy regimens with or without addition of TBI all yielded similar results, with complete responses (CR) of around 24% in both groups [9]. In contrast, mice given increasing TBI concentrations and then infused with antitumor CD8^+^ T cells experienced improved long-term survival, particularly when the mouse was given one myeloablative dose of TBI of 9Gy with stem-cell support compared to its delivery in the fractionated format [33]. These discrepancies additionally highlight the necessary investigation into the impact of these different preconditioning methods on different forms of ACT.

Overall, our findings have important implications. We demonstrate that chemotherapy and radiation-based lymphodepletion methods are not equal and have different effects on cytokine induction and engraftment of CD4^+^ T cells. We find that antitumor responses vary between these methods and suggest that we should consider supplementing this regimen to improve novel CD4 T cell therapies in patients given CTX. Supplemental therapies with ACT could include agents that would increase neutrophils or reduce the number of host cells that act as suppressor cells, perhaps by co-administering cancer vaccines and/or checkpoint blockade therapy. Overall, our data reveal new mechanisms underlying how the host immune system can be altered to augment potent Th17-based immunotherapies for cancer.

## Methods

### Mice and tumor lines

These studies were conducted using C57BL/6 and Tyrosinase-related protein 1 (TRP-1) TCR transgenic mice (Rag^-^/^-^ BWTRP-1 TCR) purchased from Jackson laboratories. The TRP-1 TCR transgenic mice (Rag^-^/^-^ BWTRP-1 TCR) are bred in house at Emory University, while C57BL/6 were purchased from the Jackson Laboratory, and tumor experiments are conducted with mice ages 6–10 weeks. Two independent mouse experiments were performed, with experimental groups (TBI + ACT and CTX + ACT) containing 10 mice per group in replicate 1 and 6 mice per group in replicate 2. Mice were randomized prior to treatment with no exclusion/inclusion criteria set. Randomization was performed by grouping mice based on small, medium, or large tumor size on Day 6 post tumor injection, and then randomly assigning an equal number of each to the various treatment groups. Treatments and tumor measurements were always performed on the same day and relative time to avoid potential effects of confounders and individuals were blinded to the treatment group of each mouse. Institutional Animal Care and Use Committee at Emory University approved the animal work, and we additionally have the support of Emory’s Division of Laboratory and Animal Resources. Initial experiments were also previously conducted at MUSC with support from Institutional Animal Care and Use Committee at MUSC and support of MUSC’s Division of Laboratory and Animal Resources. B16F10 tumors were obtained from the laboratory of Dr. Nicholas Restifo and were validated and confirmed pathogen and Mycoplasma free via PCR screen.

### T-cell cultures

#### TRP-1 cells

Transgenic TRP-1 T cells were cultured in Complete Media (CM) using Th17 polarizing conditions (IL6, IL-21, IL1b, TGFb, aIFNg, aIL4, aIL2) as described previously[31]. They were activated with 1 umol/L TRP-1106-130 peptide (SGHNCGTCRPGWRGAACNQKILTVR) in the presence of splenocytes that were irradiated at 10Gy in a 1:5 cell to splenocyte ratio and grown at a concentration of 1-1.5 x 10^6^ cells/mL. 20 ng/mL of IL23 (BioLegend) was added to the culture on Days 2 and 3, and on Day 4 cells were washed and resuspended in sterile PBS for adoptive cellular therapy (ACT) at a concentration of either 800,000 cells in 200 uL PBS per mouse or 400,000 cells in 200 uL PBS per mouse. For Th1 cells displayed in Figure 1, TRP-1 T cells were cultured in the presence of hIL-12 (3 ng/mL), aIL-4 (10 ug/mL), or hIL-2 (100 IU/mL) for 1 week, splitting as needed with the addition of 100 IU/mL of IL-2 each time cells were split. Th17 cells displayed in Figure 1 were cultured using same conditions described above but for 1 week instead of 4 days.

### Adoptive Cell Transfer (ACT)

Previously, C57BL/6 were given subcutaneous B16F10 melanoma tumors by injection of 200 uL of 5 x 10^5^ B16F10 cells in sterile PBS. Tumors were grown for 1 week prior to ACT. One day prior to ACT, mice received either 5 Gy of total body irradiation (TBI) or 200 mg/kg (4 mg/mouse) of cyclophosphamide or no preconditioning. ACT of the Th17 polarized TRP-1 cells as described above was performed by tail vein injection one week after tumor injection. .

### Cytokine multiplex assay

Serum from mice was collected on Day 10 and was then stored at −80°C prior to analysis using the Eve Technologies 44-plex Mouse Cytokine Array Discovery Assay.

### Tissue Collection and Processing

On Day 7, peripheral blood was collected from the mandibular vein into 0.125 mol/L EDTA, spun down, and resuspended in red blood cell lysis buffer (Biolegend) for 5 minutes, and then assayed using flow cytometry using the antibodies depicted in Supplemental Table 1. Peripheral blood was collected again on Day 10 and was spun down to collect serum that was sent for cytokine multiplex array (see above). On Day 39, the lymph nodes, spleen, skin, and blood were collected from the mice after euthanasia. Lymph nodes were collected into CM media and then processed into single cell suspension by mechanical dissociation over a 40 μm filter and assayed using flow cytometry. Spleens were also processed into single cell suspension by mechanical dissociation over a 40 μm filter, then resuspended in red blood cell lysis buffer (Biolegend) for 5 minutes before assaying via flow cytometry. Blood was processed as described above and similarly assayed using flow cytometry. Skin was minced and incubated in buffer containing 3 mg/mL collagenase IV (Worthington Biochemical), and 0.2 mg/mL DNase (Sigma) in Hank’s balanced salt solution at 37° C for 45 minutes with stirring. Digestion was neutralized with RPMI containing 10% FBS and 10 mmol/L EDTA. Digested and processed tissue was filtered prior to assay using flow cytometry.

### Flow cytometry

Flow cytometry was performed using BD FACSymphony™ A3 Cell Analyzer and analyzed using FlowJo software (BD Biosciences). For extracellular antibodies, the samples were resuspended in FACS buffer (PBS with 2% FBS) and incubated in antibodies at a 1:500 dilution for 20 minutes. Intracellular staining was conducted using the FoxP3/Transcription factor kit according to manufacturer’s instructions and a 1:200 dilution of antibodies (eBioscience). Antibodies used are displayed in Supplementary Tables 1 and 2.

### Statistical analysis

Comparisons of cytokines was performed using one-way ANOVA. Day 7 engraftment data was compared using one-way ANOVA. Day 39 engraftment in all organs and Ki-67 levels were compared between CTX and ACT versus TBI and ACT groups using the Mann Whitney U Test. P values are depicted numerically on the figure with values less than 0.05 considered significant. For the IFN-g and IL-17 comparison in Figure 1, a t-test was used. Kaplan–Meier survival curves were compared between treatment group pairs using the log-rank test.

## Data and materials availability

All data are available upon reasonable request.

## Declarations

Ethics approval and consent to participate: This study does not involve human participants. All mouse studies were conducted with the support of the Division of Animal Resources (DAR) and Institutional Animal Care and Use Committee under protocol: PROTO201900225 (Emory University) and 0488 (Medical University of South Carolina).

Consent for publication: Not applicable

Availability of data and material: All data associated with this study is either present in the article or will be made available upon reasonable request.

### Competing interests

The authors disclose no conflicts of interest in relation to the published work. CMP have previous received funds for consultancies/advisory boards/research contracts: Ares Immunotherapy, Lycera, Obsidian and TheromoFisher. GBL has received research funding through a sponsored research agreement between Emory University and Merck and Co., Bristol-Myers Squibb, Boerhinger-Ingelheim, and Vaccinex.

### Funding

This work was supported by the Melanoma Research Foundation (to G.O Rangel Rivera, H.M. Knochelmann, and M.C. Wittling), ARCs Foundation (to A.C. Cole), NCI F30CA243307 (HMK), NIH DE017551 (HMK), NIH R50CA233186 (to M.M. Wyatt); NCI R01CA228406, R21CA266088-01, R21CA270903 (to G.B. Lesinski), NCI R01CA175061, R01CA208514, R01CA275199 plus MUSC and Emory University Start Up Funds (to C.M. Paulos).

### Authors’ contributions

MCW and HMK: designed and conducted experiments, analyzed and interpreted data, wrote and assembled manuscript. CMP and GLB reviewed data, edited/wrote the manuscript, and helped design experiments. ACC, MMW, GORR helped to perform experiments and contributed to figures and/or text in the manuscript. Acknowledgements: We would like to acknowledge the cores both at Medical University of South Carolina and the Winship Cancer Institute/Emory University that made this research possible including the Pediatric/Winship Flow Cytometry Shared Resource, Winship Cancer Animal Models Shared Resource (NIH/NCI award number P30CA138292), and FACS Shared Resource at Hollings Cancer Center, Medical University of South Carolina (P30CA138313).

Authors’ information (optional): Megen C. Wittling ORCID https://orcid.org/0000-0002-3631-4955, Hannah M. Knochelmann https://orcid.org/0000-0003-0644-123X, Gregory B. Lesinski https://orcid.org/0000-0002-8787-7678, Chrystal M. Paulos https://orcid.org/0000-0002-0784-2601

## List of Abbreviations

Th17: T helper 17
ACT: Adoptive Cellular Therapy
TBI: Total Body Irradiation
CTX: Chemotherapy
CAR: Chimeric Antigen Receptor
G-CSF: Granulocyte colony stimulating factor
EPO: erythropoietin
IL-6: Interleukin-6
MCP-1: Monocyte Chemoattractant Protein-1
IL-5: Interleukin-5
KC: Keratinocyte chemoattractant
TIL: tumor infiltrating lymphocytes
TCR: T cell receptor
MHC: major histocompatibility complex
NK: natural killer
DC: dendritic cell
FLU: fludarabine
DAMPs: damage-associated molecular patterns
HMGB1: high-mobility group box 1 protein
IL-17: Interleukin-17
IFN: interferon
TRP: tyrosinase-related peptide
Th1: T helper 1

## Supplemental Materials

**Supplemental Table 1:** Flow Panel Used for Flow Cytometry on whole blood collected Day 7 after ACT.

| Species | Color | Marker | Catalog Number | Clone | Vendor |
| --- | --- | --- | --- | --- | --- |
| Mouse | BUV395 | CD3e | 740268 | 17A2 | BD Biosciences |
| Mouse | BUV737 | CD8a | 612759 | 53-6.7 | BD Biosciences |
| NA | BUV459 | Zombie UV | 423108 | NA | BioLegend |
| Mouse | BV750 | CD4 | 100467 | GK1.5 | Biolegend |
| Mouse | APC Cy7 | CD90.1 (Thy1.1) | 202520 | OX-7 | Biolegend |
| Mouse | BV786 | CD90.2 (Thy1.2) | 564365 | 53-2.1 | BD Biosciences |
| Mouse | FITC | V β14 | 553258 | 14-2 | BD Biosciences |
| Mouse | BV650 | Ki67 | 151215 | 11F6 | BioLegend |

**Supplemental Table 2:** Flow Panel Used for Flow Cytometry on Skin, Lymph nodes, and spleen tissues collected at endpoint (day 39)

| Species | Color | Marker | Catalog Number | Clone | Vendor |
| --- | --- | --- | --- | --- | --- |
| Mouse | BUV395 | CD3e | 740268 | 17A2 | BD Biosciences |
| NA | BUV 459 | Zombie UV (LD) | 423108 | NA | BioLegend |
| Mouse | BUV737 | CD8a | 612759 | 53-6.7 | BD Biosciences |
| Mouse | BV650 | Ki67 | 151215 | 11F6 | BioLegend |
| Mouse | BV750 | CD4 | 100467 | GK1.5 | BioLegend |
| Mouse | FITC | V $\beta$ 14 | 553258 | 14-2 | BD Biosciences |

## Notes

**Conflict of Interest**: The authors declare no conflict of interest.

### Competing Interest Statement

The authors have declared no competing interest.

## References

1. Giordano Attianese, G.M.P., S. Ash, and M. Irving, Coengineering specificity, safety, and function into T cells for cancer immunotherapy. Immunol Rev, 2023.

2. Knochelmann, H.M., et al., CAR T Cells in Solid Tumors: Blueprints for Building Effective Therapies. Front Immunol, 2018. 9: p. 1740.

3. Hossian, A., et al., Multipurposing CARs: Same engine, different vehicles. Mol Ther, 2022. 30(4): p. 1381–1395.

4. Akce, M., et al., The Potential of CAR T Cell Therapy in Pancreatic Cancer. Front Immunol, 2018. 9: p. 2166.

5. Rosenberg, S.A. and M.E. Dudley, Adoptive cell therapy for the treatment of patients with metastatic melanoma. Curr Opin Immunol, 2009. 21(2): p. 233–40.

6. Rohaan, M.W., S. Wilgenhof, and J. Haanen, Adoptive cellular therapies: the current landscape. Virchows Arch, 2019. 474(4): p. 449–461.

7. Porter, D.L., et al., Chimeric antigen receptor-modified T cells in chronic lymphoid leukemia. N Engl J Med, 2011. 365(8): p. 725–33.

8. Maude, S.L., et al., Chimeric antigen receptor T cells for sustained remissions in leukemia. N Engl J Med, 2014. 371(16): p. 1507–17.

9. Goff, S.L., et al., Randomized, Prospective Evaluation Comparing Intensity of Lymphodepletion Before Adoptive Transfer of Tumor-Infiltrating Lymphocytes for Patients With Metastatic Melanoma. J Clin Oncol, 2016. 34(20): p. 2389–97.

10. Leidner, R., et al., Neoantigen T-Cell Receptor Gene Therapy in Pancreatic Cancer. N Engl J Med, 2022. 386(22): p. 2112–2119.

11. Paulos, C.M., et al., Toll-like receptors in tumor immunotherapy. Clin Cancer Res, 2007. 13(18 Pt 1): p. 5280–9.

12. Gattinoni, L., et al., Adoptive immunotherapy for cancer: building on success. Nat Rev Immunol, 2006. 6(5): p. 383–93.

13. Salem, M.L., et al., Defining the ability of cyclophosphamide preconditioning to enhance the antigen-specific CD8+ T-cell response to peptide vaccination: creation of a beneficial host microenvironment involving type I IFNs and myeloid cells. J Immunother, 2007. 30(1): p. 40–53.

14. Paulos, C.M., et al., Microbial translocation augments the function of adoptively transferred self/tumor-specific CD8+ T cells via TLR4 signaling. J Clin Invest, 2007. 117(8): p. 2197–204.

15. Oh, D.Y. and L. Fong, Cytotoxic CD4(+) T cells in cancer: Expanding the immune effector toolbox. Immunity, 2021. 54(12): p. 2701–2711.

16. Nelson, M.H., et al., Identification of human CD4(+) T cell populations with distinct antitumor activity. Sci Adv, 2020. 6(27).

17. Johnson, C.B., et al., Enhanced Lymphodepletion Is Insufficient to Replace Exogenous IL2 or IL15 Therapy in Augmenting the Efficacy of Adoptively Transferred Effector CD8(+) T Cells. Cancer Res, 2018. 78(11): p. 3067–3074.

18. Bronte, V., et al., Apoptotic death of CD8+ T lymphocytes after immunization: induction of a suppressive population of Mac-1+/Gr-1+ cells. J Immunol, 1998. 161(10): p. 5313–20.

19. Antony, P.A., et al., CD8+ T cell immunity against a tumor/self-antigen is augmented by CD4+ T helper cells and hindered by naturally occurring T regulatory cells. J Immunol, 2005. 174(5): p. 2591–601.

20. Turk, M.J., et al., Concomitant tumor immunity to a poorly immunogenic melanoma is prevented by regulatory T cells. J Exp Med, 2004. 200(6): p. 771–82.

21. Wallen, H., et al., Fludarabine modulates immune response and extends in vivo survival of adoptively transferred CD8 T cells in patients with metastatic melanoma. PLoS One, 2009. 4(3): p. e4749.

22. Gattinoni, L., et al., Removal of homeostatic cytokine sinks by lymphodepletion enhances the efficacy of adoptively transferred tumor-specific CD8+ T cells. J Exp Med, 2005. 202(7): p. 907–12.

23. Citrin, D.E., Recent Developments in Radiotherapy. N Engl J Med, 2017. 377(22): p. 2200–2201.

24. Viaud, S., et al., Cyclophosphamide induces differentiation of Th17 cells in cancer patients. Cancer Res, 2011. 71(3): p. 661–5.

25. Schiavoni, G., et al., Cyclophosphamide induces type I interferon and augments the number of CD44(hi) T lymphocytes in mice: implications for strategies of chemoimmunotherapy of cancer. Blood, 2000. 95(6): p. 2024–30.

26. Di Roio, A., et al., Mdr1-Expressing Cd4(+) T Cells with Th1.17 Features Resist to Neoadjuvant Chemotherapy and Are Associated with Breast Cancer Clinical Response. J Immunother Cancer, 2023. 11(11).

27. Hanada, K.I., et al., An effective mouse model for adoptive cancer immunotherapy targeting neoantigens. JCI Insight, 2019. 4(10).

28. Ward-Kavanagh, L.K., et al., Whole-body irradiation increases the magnitude and persistence of adoptively transferred T cells associated with tumor regression in a mouse model of prostate cancer. Cancer Immunol Res, 2014. 2(8): p. 777–88.

29. Zhu, E.F., et al., Synergistic innate and adaptive immune response to combination immunotherapy with anti-tumor antigen antibodies and extended serum half-life IL-2. Cancer Cell, 2015. 27(4): p. 489–501.

30. Xie, Y., et al., Naive tumor-specific CD4(+) T cells differentiated in vivo eradicate established melanoma. J Exp Med, 2010. 207(3): p. 651–67.

31. Knochelmann, H.M., et al., IL6 Fuels Durable Memory for Th17 Cell-Mediated Responses to Tumors. Cancer Res, 2020. 80(18): p. 3920–3932.

32. Dudley, M.E., et al., Adoptive cell therapy for patients with metastatic melanoma: evaluation of intensive myeloablative chemoradiation preparative regimens. J Clin Oncol, 2008. 26(32): p. 5233–9.

33. Wrzesinski, C., et al., Increased intensity lymphodepletion enhances tumor treatment efficacy of adoptively transferred tumor-specific T cells. J Immunother, 2010. 33(1): p. 1–7.

34. Dudley, M.E., et al., Adoptive cell transfer therapy following non-myeloablative but lymphodepleting chemotherapy for the treatment of patients with refractory metastatic melanoma. J Clin Oncol, 2005. 23(10): p. 2346–57.

35. Bechman, N. and J. Maher, Lymphodepletion strategies to potentiate adoptive T-cell immunotherapy - what are we doing; where are we going? Expert Opin Biol Ther, 2021. 21(5): p. 627–637.

36. Paulos, C.M., et al., The inducible costimulator (ICOS) is critical for the development of human T(H)17 cells. Sci Transl Med, 2010. 2(55): p. 55ra78.

37. Bailey, S.R., et al., Th17 cells in cancer: the ultimate identity crisis. Front Immunol, 2014. 5: p. 276.

38. Muranski, P., et al., Tumor-specific Th17-polarized cells eradicate large established melanoma. Blood, 2008. 112(2): p. 362–73.

39. Lai, P., et al., C3aR costimulation enhances the antitumor efficacy of CAR-T cell therapy through Th17 expansion and memory T cell induction. J Hematol Oncol, 2022. 15(1): p. 68.

40. Guedan, S., et al., Single residue in CD28-costimulated CAR-T cells limits long-term persistence and antitumor durability. J Clin Invest, 2020. 130(6): p. 3087–3097.

41. Bowers, J.S., et al., Th17 cells are refractory to senescence and retain robust antitumor activity after long-term ex vivo expansion. JCI Insight, 2017. 2(5): p. e90772.

42. Neitzke, D.J., et al., Murine Th17 cells utilize IL-2 receptor gamma chain cytokines but are resistant to cytokine withdrawal-induced apoptosis. Cancer Immunol Immunother, 2017. 66(6): p. 737–751.

43. Kryczek, I., et al., Human TH17 cells are long-lived effector memory cells. Sci Transl Med, 2011. 3(104): p. 104ra100.

44. Kalos, M., et al., T cells with chimeric antigen receptors have potent antitumor effects and can establish memory in patients with advanced leukemia. Sci Transl Med, 2011. 3(95): p. 95ra73.

45. Yang, J.C. and S.A. Rosenberg, Adoptive T-Cell Therapy for Cancer. Adv Immunol, 2016. 130: p. 279–94.

46. Wagner, J., et al., CAR T Cell Therapy for Solid Tumors: Bright Future or Dark Reality? Mol Ther, 2020. 28(11): p. 2320–2339.

47. Kirtane, K., et al., Adoptive cellular therapy in solid tumor malignancies: review of the literature and challenges ahead. J Immunother Cancer, 2021. 9(7).

48. Wang, E., et al., Improving the therapeutic index in adoptive cell therapy: key factors that impact efficacy. J Immunother Cancer, 2020. 8(2).

49. Liu, Z., et al., Immunosuppression in tumor immune microenvironment and its optimization from CAR-T cell therapy. Theranostics, 2022. 12(14): p. 6273–6290.

50. Klebanoff, C.A., et al., Sinks, suppressors and antigen presenters: how lymphodepletion enhances T cell-mediated tumor immunotherapy. Trends Immunol, 2005. 26(2): p. 111–7.

51. Cachot, A., et al., Tumor-specific cytolytic CD4 T cells mediate immunity against human cancer. Sci Adv, 2021. 7(9).

52. Kamphorst, A.O. and R. Ahmed, CD4 T-cell immunotherapy for chronic viral infections and cancer. Immunotherapy, 2013. 5(9): p. 975–87.

53. Bailey, S.R., et al., Human CD26(high) T cells elicit tumor immunity against multiple malignancies via enhanced migration and persistence. Nat Commun, 2017. 8(1): p. 1961.

54. De Filippo, K., et al., Mast cell and macrophage chemokines CXCL1/CXCL2 control the early stage of neutrophil recruitment during tissue inflammation. Blood, 2013. 121(24): p. 4930–7.

55. Theyab, A., et al., New insight into the mechanism of granulocyte colony-stimulating factor (G-CSF) that induces the mobilization of neutrophils. Hematology, 2021. 26(1): p. 628–636.

56. Leitch, V.D., et al., IL-5-overexpressing mice exhibit eosinophilia and altered wound healing through mechanisms involving prolonged inflammation. Immunol Cell Biol, 2009. 87(2): p. 131–40.

57. Fisher, J.W., Landmark advances in the development of erythropoietin. Exp Biol Med (Maywood), 2010. 235(12): p. 1398–411.

58. Hirschhorn, D., et al., T cell immunotherapies engage neutrophils to eliminate tumor antigen escape variants. Cell, 2023. 186(7): p. 1432–1447 e17.

59. Rosenberg, S.A., et al., Durable complete responses in heavily pretreated patients with metastatic melanoma using T-cell transfer immunotherapy. Clin Cancer Res, 2011. 17(13): p. 4550–7.

